# Cell-based simulations of Notch-dependent cell differentiation on growing domains

**DOI:** 10.1101/859363

**Authors:** Anna Stopka, Marcelo Boareto, Dagmar Iber

## Abstract

Notch signalling controls cell differentiation and proliferation in many tissues. The Notch signal is generated by the interaction between the Notch receptor of one cell with the Notch ligand (Delta or Jagged) of a neighbouring cell. Therefore, the pathway requires cell-cell contact in order to be active. During organ development, cell differentiation occurs concurrently with tissue growth and changes in cell morphology. How growth impacts on Notch signalling and cell differentiation remains poorly understood. Here, we developed a modelling environment to simulate Notch signalling in a growing tissue. We use our model to simulate the differentiation process of pancreatic progenitor cells. Our results suggest that Notch-mediated differentiation in the developing pancreas is first mediated by geometric effects that result in loss of Notch signalling on the tissue boundary, leading to the differentiation of tip versus trunk cells. A second wave of differentiation further happens in the trunk cells due to a reduction in the expression of the ligand *Jagged*, which has been shown to be controlled by signalling factors secreted from the surrounding mesenchyme. Our results bring new insights into how cells coordinate tissue growth with cell fate specification during organ development.

## Introduction

The pancreas develops from an epithelial bud that evaginates from the foregut into the surrounding mesenchyme. After an initial expansion step, the pancreatic progenitor cells at the surface of the epithelium adopt a tip identity, while the progenitor cells in the interior of the tissue adopt a trunk identity (Shih et al 2013). The tip/trunk separation is mediated by Notch signalling, where Notch activity promotes trunk and represses tip identity (Kim et al, 2010). Once the tip/trunk decision is established, the outer tip layer starts to branch, while trunk cells differentiate into either ductal or endocrine cells (Li et al, 2015). This second cell fate decision is also mediated by Notch-signalling, where high Notch activity favours the ductal fate. In addition to Notch, FGF signalling plays a critical role in pancreas development. FGF10 signalling is required for proper cell proliferation (Bhushan et al, 2001; Dichmann et al, 2003; Seymour et al, 2012). In *Fgf10* mutant embryos, the initial evagination of the pancreatic epithelium is normal, but subsequent growth is impaired (Bhushan et al, 2001). Consistently, persistent expression of *Fgf10* stimulates cell proliferation, and also inhibits differentiation by maintaining Notch activity (Hart et al, 2003). The effect of FGF10 on cell differentiation has been shown to be mediated by Notch signalling, likely via the upregulation of the expression of the Notch ligand *Jagged* (Norgaard et al, 2003). Although many key signalling effectors and transcription factors that regulate pancreatic differentiation have been identified, there is little understanding about their interactions and how the cells interpret different signals as the tissue grows during pancreas development.

The Notch signalling pathway is active when the Notch receptor of one cell interacts with the ligand (Delta or Jagged) of a neighbouring cell, releasing the Notch Intracellular Domain (NICD) (Artavanis-Tsakonas et al, 1999; Bray, 2016; Andersson et al, 2011). Interestingly, the same signal (NICD) is triggered by either Delta or Jagged ligands. However, due to differences in how the expression of these ligands is regulated by NICD, different tissue patterns are observed, depending on which ligand is more abundant (Boareto et al, 2015; Sjöqvist and Andersson, 2017). Delta (DLL1) is negatively modulated by NICD (Heitzler et al, 1996; Artavanis-Tsakonas et al, 1999), leading to an intercellular negative feedback loop that drives neighbouring cells to adopt alternate states: one cell keeps high Notch signal and low Delta (Receiver cell), while the neighbouring cell adopts an alternate state by having low Notch signal and high Delta (Sender cell) (Collier et al, 1996; Sprinzak et al, 2010). This mechanism is referred to as lateral inhibition and has been shown to control cell differentiation in many tissues (de la Pompa et al, 1997). In contrast to DLL1, *Jagged1* expression is usually activated by NICD, which leads to cell fate propagation in many organs, in a process referred to as lateral induction (Ross and Kadesch, 2004; Manderfield et al, 2012; Petrovic et al, 2014).

In a growing tissue, there are frequent cell rearrangements. This can lead to dynamic changes in cell morphology, such as cell shape, contact area, and cell neighbours. Notch signalling activity has been shown to be highly dependent on cell morphological properties such as cell-cell adhesion (Hatakeyama et al, 2014), cell-cell contact area (Shaya et al, 2017), and asymmetry in cell division (Kosodo et al, 2004). Here, to facilitate the study of the effect of cell morphology and neighbour relationships on Notch activity, we established a 2D cell-based simulation environment to simulate Notch-mediated cell differentiation in a growing epithelial tissue. To this end, we incorporated Notch interactions into the 2D cell-based environment (`LBIBCell`) (Tanaka et al, 2015). `LBIBCell` has many similarities with vertex models (Fletcher et al, 2014), but there are important differences. In `LBIBCell`, each cell is a separate geometric entity and the high geometric resolution permits curved boundaries between cells. Cells interact with their neighbours via spring-like cell-cell junctions. Moreover, `LBIBCell` explicitly represents the fluid inside (cytoplasm) and outside (interstitial fluid) the cells using the Lattice Boltzmann (LB) method (Chen et al, 1998). The Immersed Boundary (IB) method is used to model the fluid-structure interaction (Peskin et al, 2002). Growth of the cells is simulated by adding mass to intracellular grid points and cells that exceed a certain cell area are divided. We have previously defined the parameter ranges that allow us to recapitulate epithelial tissue dynamics (Kokic et al, 2019; Vetter et al 2019; Stopka et al 2019). The set of coupled ordinary differential equations (ODEs) describing the Notch signalling network (Sprinzak et al, 2010; Boareto et al, 2015) can be solved on the cell boundary grid points, using the LB method.

Using our simulation environment, we explored the impact of the growth rate, of asymmetric cell division, and of the expression of Notch ligands on cell differentiation. We find that geometric effects can result in inactive Notch signalling and thus cell differentiation on the tissue boundary. This effect can potentially explain why future tip cells first emerge on the outer boundary of the pancreatic bud. We also find that the later differentiation of trunk cells can, in principle, be explained with an FGF10-dependent reduction in *Jagged* expression.

## Results

### Effect of asymmetry in cell division on Notch-mediated cell differentiation

Previous theoretical efforts have shown that in the presence of high levels of the ligand Jagged, all cells adopt similar levels of Notch activity and no pattern emerges (Boareto et al, 2015; Jolly et al, 2015). In contrast, when Jagged levels are low or absent, Notch-Delta interactions predominate and the cells tend to alternate their state, forming checkerboard spatial pattern (Collier et al, 1996; Sprinzak et al, 2010). For this reason, we used low levels of Jagged in our simulations to study the effect of asymmetry in cell division on Notch-mediated patterning. In our simulations, each tissue was comprised of about 400 cells that were initially undifferentiated, i.e., the cells start with high levels of Notch activity.

We observed that the cells form a checkerboard spatial pattern, typical for Notch-Delta interactions, for different levels of asymmetry during cell division. Moreover, as the asymmetry in cell division increases the tissue becomes more heterogeneous both in cell size and in the levels of Notch signalling (Figure 1a-d). We also observed that cells with low Notch signal, and consequently high levels of Delta, tend to be smaller (Figure 1a,c). This is consistent with experimental observations showing that the cell contact area affects Notch signalling and that smaller cells are more likely to differentiate, as observed in hair cell precursors in the inner ear (Shaya et al, 2017).

We further evaluated whether different levels of asymmetry in cell division impacts on cell differentiation. For that, we assume that once a cell reaches a low level of Notch signal it differentiates and remains differentiated even if the Notch signal levels increase afterwards. We also assume that differentiated cells proliferate with the same rate as undifferentiated cells. Under these conditions, we found that asymmetric cell division does not impact on the differentiation rate (Figure 1f,e).

**Figure 1.**
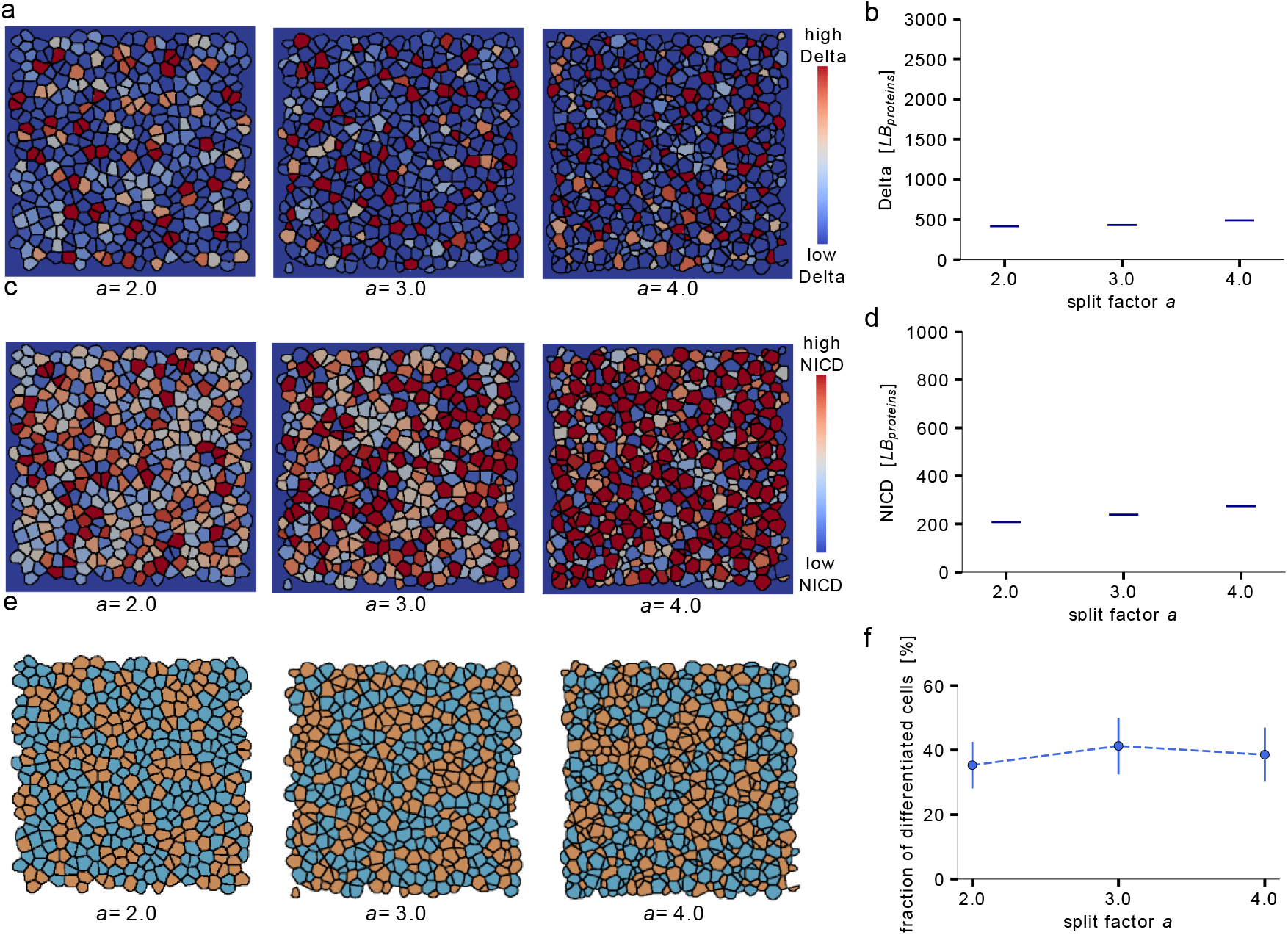
Asymmetry in cell divisions affects Notch-mediated patterning but does not affect cell differentiation rate. (a) Levels of Delta for a tissue with symmetric cell divisions and asymmetric cell divisions. Asymmetry is quantified by the asymmetry factor α, such that (α = 2) for symmetric cell divisions and (α> 2) for asymmetric cell divisions. Asymmetry in cell division increases with the increase of α (Methods). (b) Asymmetry in cell division has little impact on the levels of Delta. (c) Levels of Notch signal (NICD) for a tissue with symmetric cell divisions (α = 2) and asymmetric cell divisions (α = 3, α = 4). (d) Asymmetry in cell division leads to a broader distribution of NICD levels, but has little impact on its average levels. (e) A qualitative comparison of cell fate patterns for symmetric and asymmetric divisions shows no major pattern difference. We considered that a cell differentiates when it has low Notch signal (NICD < 100) (Methods). Differentiated cells (orange) have the same proliferation rate as undifferentiated cells (blue). (f) The fraction of differentiated cells is around 40% for different levels of asymmetry in cell division.

### Impact of the growth rate on Notch-mediated cell differentiation

We further studied the impact of the growth rate on Notch signalling dynamics. To do so, we simulated tissue growth for different growth rates with symmetric cell division. Once again, we observed the typical salt and pepper patterning for different growth rates (Figure 2a). Interestingly, we observed that Notch activity becomes more homogeneous in a fast-growing tissue. Fewer cells reach high levels of Delta (Figure 2b), and the cells tend to have similar levels of Notch signal (Figure 2c,d). Also, the number of cells with low Notch activity decreases. These results suggest that fast growth leads to a dynamic change in the neighbours that can lead to the maintenance of Notch activity in most cells (Figure 2d).

**Figure 2.**
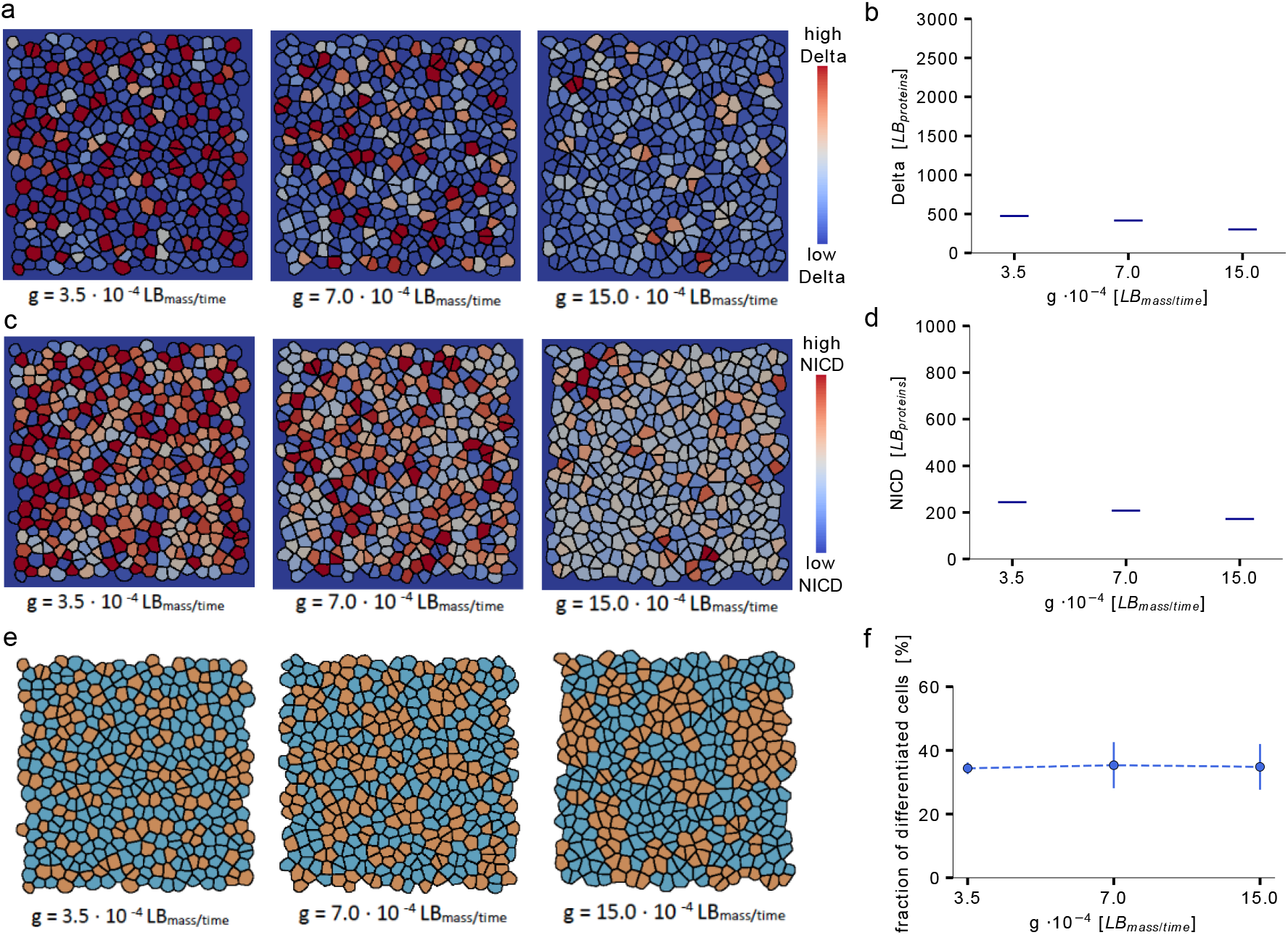
Growth rate strongly affects Notch signal distribution, but has little impact on cell differentiation rate. (a) Levels of Delta for a tissue with different growth rates. A growth source of g=(3.5, 7.0, 15.0) 10_-4_ LB_mass/time_ corresponds to an average cell cycle of approximately (48, 24, 12) hours, respectively. (b) As the growth rate increases less cells adopt high Delta levels. (c) Levels of NICD for a tissue with different growth rates. (d) The distribution of NICD levels become more homogenous and fewer cells have low NICD levels as the growth rate increases. (e) A qualitative comparison of cell fate patterns for different growth rates reveals the formation of clusters of differentiated cells in fast-growing tissues. We considered that a cell differentiates when it has low Notch signal (NICD < 100) (Methods). Differentiated cells (orange) have the same proliferation rate as undifferentiated cells (blue). (f) The fraction of differentiated cells remains the same for all growth rates.

When considering cell differentiation, we noted a significant difference in the spatial distribution of differentiated cells depending on how fast the tissue grows. As the growth rate increases, differentiated cells form bigger clusters (Figure 2e). However, we observed that such differences in the distribution of differentiated cells does not affect the differentiation rate, which remains the same for different growth rates (Figure 2f). We note that for much lower NICD threshold levels, the rate of cell differentiation would increase at lower growth rates. However, for such a low NICD threshold, the absolute rate of cell differentiation would be low. This effect is therefore unlikely to explain the increased rates of cell differentiation as growth rates decline during development.

### Tissue geometry and Jagged signalling affect cell fate pattern

We next sought to study the effect of the tissue geometry on Notch-mediated cell fate determination. To do so, we simulated a tissue growing in free space, i.e., the tissue did not get close to the boundaries of the simulated domain. The simulation started with an initial tissue structure of about 100 multipotent pancreatic cells (MPCs). As previously described, the cell differentiation in our simulations was coupled to Notch signalling: high Notch concentrations promote progenitor cell maintenance, while low Notch concentrations in a cell drive its differentiation. We considered that MPCs are of cell type 1 (blue cells) and cells with a low NICD concentration (NICD < 100) committed irreversibly to differentiation (orange cells, type 2). When dividing, cells inherited their own cell fate to their progeny. We observed that cells in the outer cell layer of a tissue have a higher chance to differentiate than cells at the inner part of a tissue (Figure 3a). Cells at the tissue boundary have fewer cell neighbours and therefore fewer signalling interactions. As a consequence, their Notch and thus NICD concentrations tend to be lower, which leads to cell differentiation.

**Figure 3.**
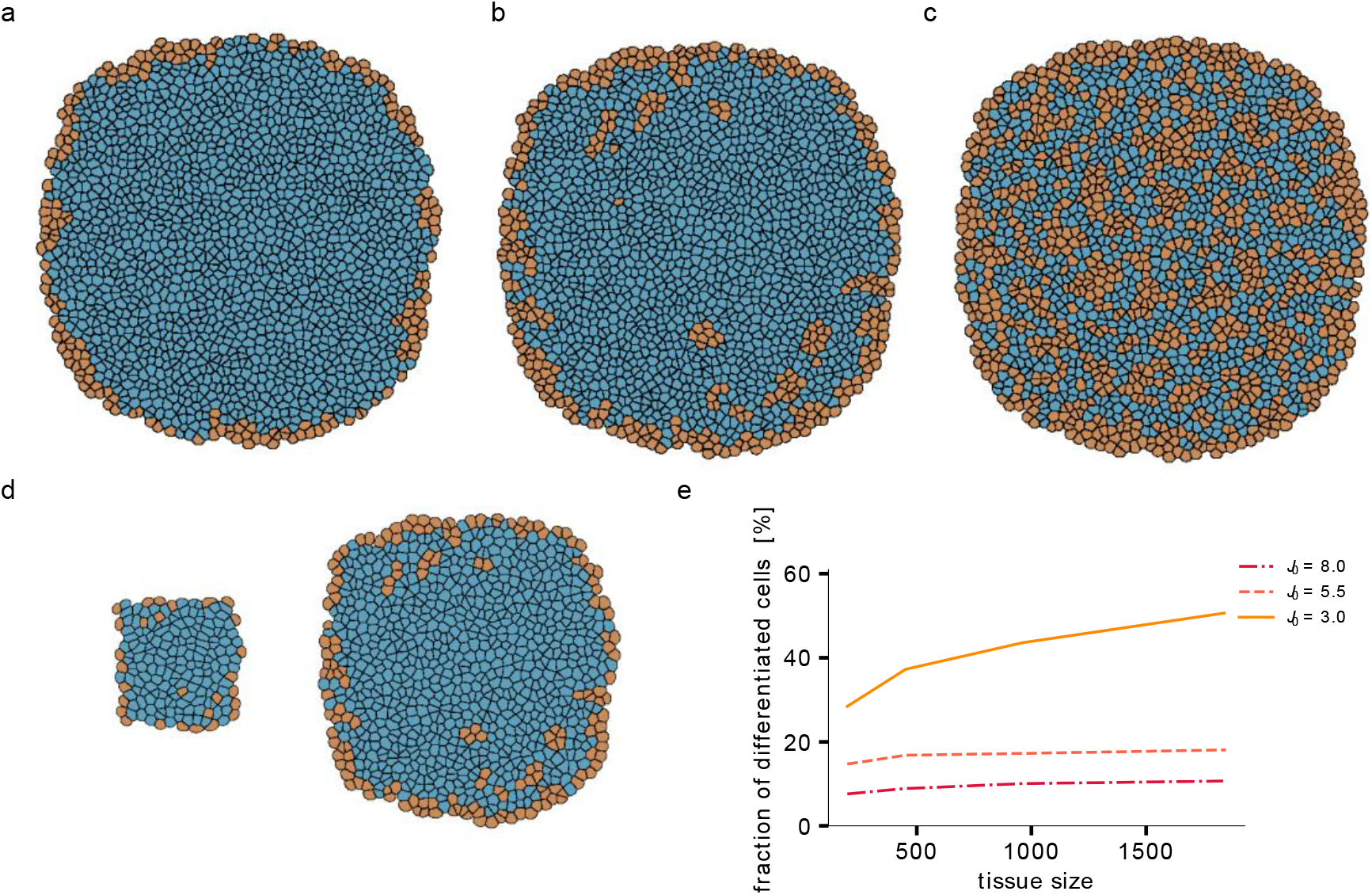
Tissue geometry and Jagged signalling affect Notch mediated cell fate determination. (a-c) Simulated tissue with high, intermediate and low levels of Jagged expression, respectively. (a) For high Jagged expression only cells at the tissue boundary differentiates. Due to a lower number of neighbours, boundary cells tend to have lower Notch concentrations, which generates a decrease in the Notch concentration driving cell differentiation. (b) Even for intermediate Jagged expression levels most of the differentiated cells are at the tissue boundary. (c) Only for low Jagged expression levels many cells at the tissue inner differentiate. (d,e) Fraction of differentiated cells scales with tissue size for high and intermediated Jagged expression levels, while for low values of Jagged expression it increases with the tissue size.

In addition, we tested the effect of signalling on the cell fate determination process in the pancreas epithelium. FGF signalling not only promotes outgrowth, but also supports *Jagged* expression (Li et al, 2016; Mitsiadis et al, 2010). We find that *Jagged* expression levels have a strong impact on cell differentiation in the centre of the tissue, but not on differentiation on the tissue boundary (Figure 3). As the levels of *Jagged* expression decrease, differentiation increases in the centre of the tissue (Figure 3a-c). In case of high Jagged production rates only cells at the boundary differentiate, and the fraction of differentiated cells is independent of the tissue size (Figure 3d,e). This suggests that the outer ring of differentiated cells is formed due to the tissue geometry (Figure 3a-c), while the inner part is under control of signalling.

In summary, our simulations revealed two findings: first, cells at the tissue boundary are more likely to differentiate, independent of Jagged signalling, and second, the fraction of differentiated cells in the inner part of the tissue depends on the Jagged signalling level.

### Tissue geometry and limited access to surrounding mesenchyme recapitulate two rounds of differentiation in the developing pancreas

We next tested, whether the two observed effects on Notch mediated cell fate determination (geometry/boundary effect and decreased Jagged levels) could explain the two rounds of differentiation (tip vs trunk and ductal vs endocrine) observed during pancreas development. For that, we simulated a growing tissue for approximately 4 days, corresponding to a growing mouse pancreas from embryonic day E11.5 to E15.5. Based on experimental observations in mouse (Kim et al, 2015), we used a cell cycle of 24 hours. Also, we simulated the decreasing influence of FGF signalling from the surrounding mesenchyme by considering that the expression levels of *Jagged* decreases over time.

In the beginning of the simulation, which corresponds to embryonic day E11.5, the tissue is comprised of undifferentiated MPCs (Figure 4a). As the simulation starts, we observed an increase in the differentiation rate until it becomes relatively constant after one day (Figure 4b). This increase in differentiation corresponds to the differentiation of cells at the boundary of the tissue (Figure 4a). A second increase in the differentiation rate occurs around embryonic day E14.5. This second wave of differentiation arises from the differentiation of the inner cells, which occurs only when *Jagged* expression levels are low (Figure 4a,b). Taken together, these results suggest that boundary effects together with decreased *Jagged* expression can lead to two distinct waves of differentiation. Interestingly, these two waves of differentiation are present but not clearly separated in a tissue without growth, where the second wave of differentiation occurs earlier (Figure 4b). This suggests that growth can delay the onset of differentiation in the inner part of the tissue, possibly by decreasing tissue heterogeneity and hampering Notch-Delta mediated cell differentiation (Figure 2c,d).

**Figure 4.**
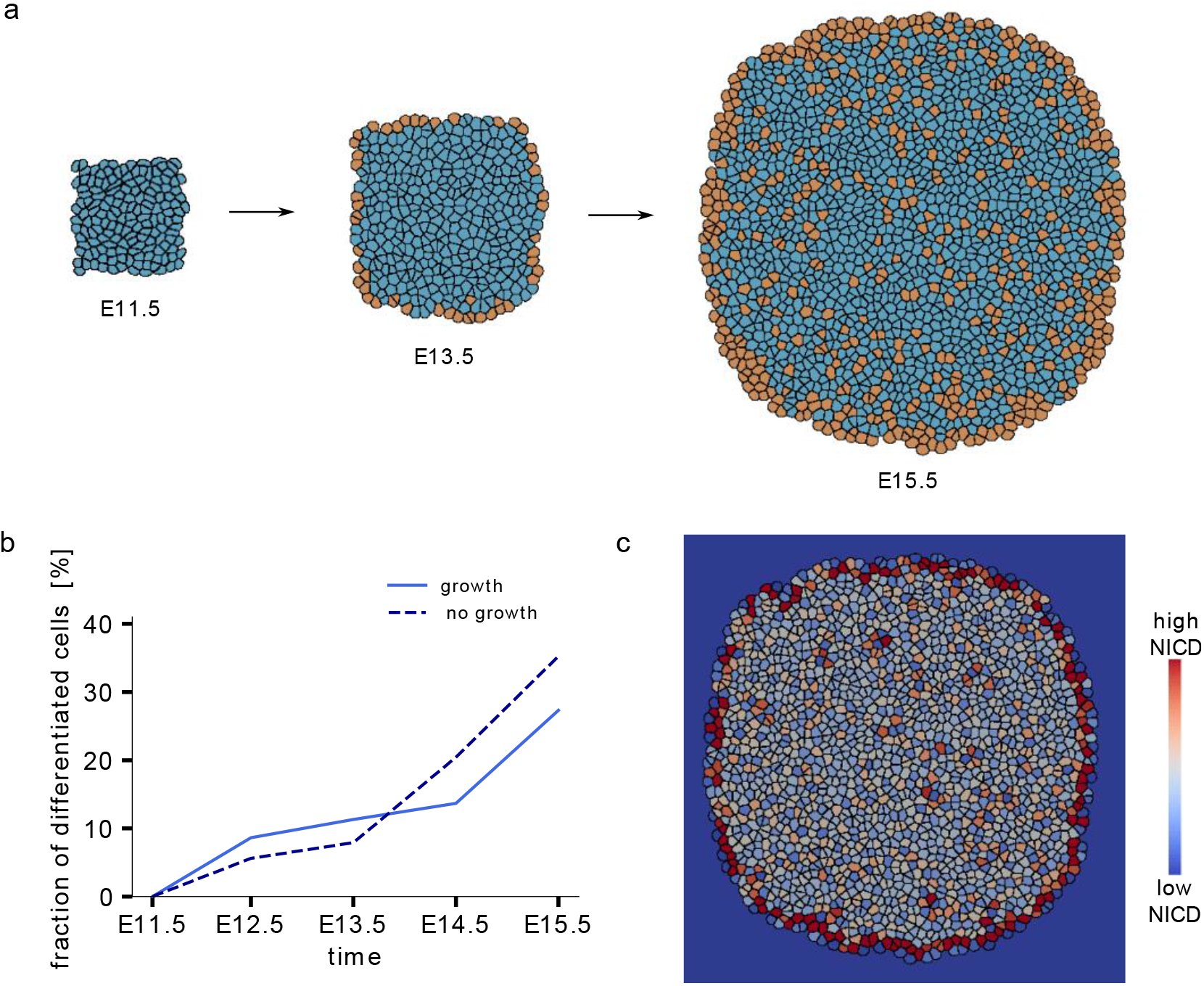
Simulating cell fate determination in the early pancreas. (a) Cell differentiation pattern for a FGF-induced linear decrease in Jagged production rates on a growing tissue at embryonic day E11.5, E13.5 and E15.5. (b) A first increase in the differentiation rates is observed due to the differentiation of the cells at the boundary of the tissue. After E12.5 differentiation rate is relatively constant and most differentiated cells are present at the boundary of the tissue. A second increase in differentiation begins at E14.5, due to the subsequent differentiation of the cells in the centre of the tissue. In the absence of growth there is no clear separation between these two differentiation waves. (c) NICD concentration patterning at E15.5: cells at the boundary (tip) and several cells at the inner of the tissue (trunk) have low NICD concentrations.

## Discussion

In the developing pancreas, several subsequent cell differentiation processes take place in order to build a complex multi-functional organ from a homogeneous tissue of undifferentiated multipotent progenitor cells (Shih et al 2013). During early pancreas development, a first wave of differentiation arises from the differentiation of progenitor cells at the boundary of the tissue into tip cells, while the cells at the centre retain the trunk identity (Kim et al, 2010). A second wave of differentiation further emerges from the differentiation of trunk cells into ductal or endocrine cell types (Li et al, 2015). Two key signalling pathways, Notch and FGF, have been shown to be pivotal during pancreas development. Notch activity has been shown to control both tip/trunk and ductal/endocrine differentiation (Shih et al 2013), while FGF activity has been largely associated to cell proliferation (Bhushan et al, 2001; Dichmann et al, 2003; Seymour et al, 2012). Recent evidence also shows that perturbations of FGF signalling affects cell differentiation, and that this effect is likely due to the FGF-dependent modulation of the expression of the Notch ligand *Jagged* (Norgaard et al, 2003).

Here, we established a simulation environment to simulate Notch-mediated cell differentiation in growing tissues. We used our simulation framework to test whether asymmetric cell divisions, growth, or a temporal decrease in *Jagged* expression can impact on Notch-mediated cell differentiation and potentially explain the differentiation processes of pancreatic progenitors. We found that asymmetry in cell division has little impact on the distribution of Notch levels or on the rate of cell differentiation. Interestingly, we noted smaller cells to be more likely to differentiate, which is consistent with previous experimental observations showing that Notch signalling is dependent on cell-cell contact area and that hair-cell precursors in the inner ear tend to be smaller (Shaya et al, 2017). We further evaluated the effect of the growth rate and observed that in fast-growing tissues, Notch activity becomes more homogeneous and fewer cells have low Notch activity. Moreover, we found that growth affects the patterning of differentiated cells. The faster the tissue grows, the bigger are the clusters of differentiated cells. Nevertheless, the differentiation rates were independent of the growth rate.

Interestingly, we found that cells at the boundary are more likely to differentiate. In the presence of high expression of *Jagged*, only cells at the boundary differentiate, and as *Jagged* expression decreases, differentiation also occurs at the centre of the tissue. By simulating a growing tissue with a temporal linear decrease in *Jagged* expression, we could recapitulate the two subsequent rounds of Notch-mediated cell differentiation observed during pancreas development. Based on these observations, it is possible that the two waves of Notch-mediated cell differentiation in the pancreas epithelium are the result of the combined effect of the tissue boundary, which leads to tip vs trunk differentiation, and a decreasing access to the mesenchymal FGF10 signalling source, which regulates *Jagged* expression leading to the differentiation of trunk cells. Whether this regulatory mechanism is sufficient to explain cell fate determination in the developing pancreas, or other mechanisms such as heterogeneous mechanical stress (Chan et al, 2017) also play a major rule remains to be determined.

## Methods

### Notch signalling in a growing tissue

In this study, we model the Notch signalling dynamics according to the model proposed by (Boareto et al, 2015). This model extends the Notch-Delta model proposed by (Sprinzak et al, 2010) by also considering the ligand Jagged (Figure 5a). In the model, Notch signalling is active when a Notch receptor of one cell interacts with the ligand of a neighbouring cell (*trans*-interaction). This leads to the release of the Notch intracellular domain (NICD) that works as a transcription factor and modulates the expression of many genes. The presence of NICD in a cell results in transcriptional activation of *Notch* and *Jagged*, and transcriptional inhibition of *Delta*. We also considered the interaction between the receptor and the ligand from the same cell (*cis*-interaction). This interaction is also known as *cis*inhibition and has been shown to be important to facilitate Notch-Delta patterning (Sprinzak et al, 2010; Formosa-Jordan and Ibañes, 2014).

**Figure 5.**
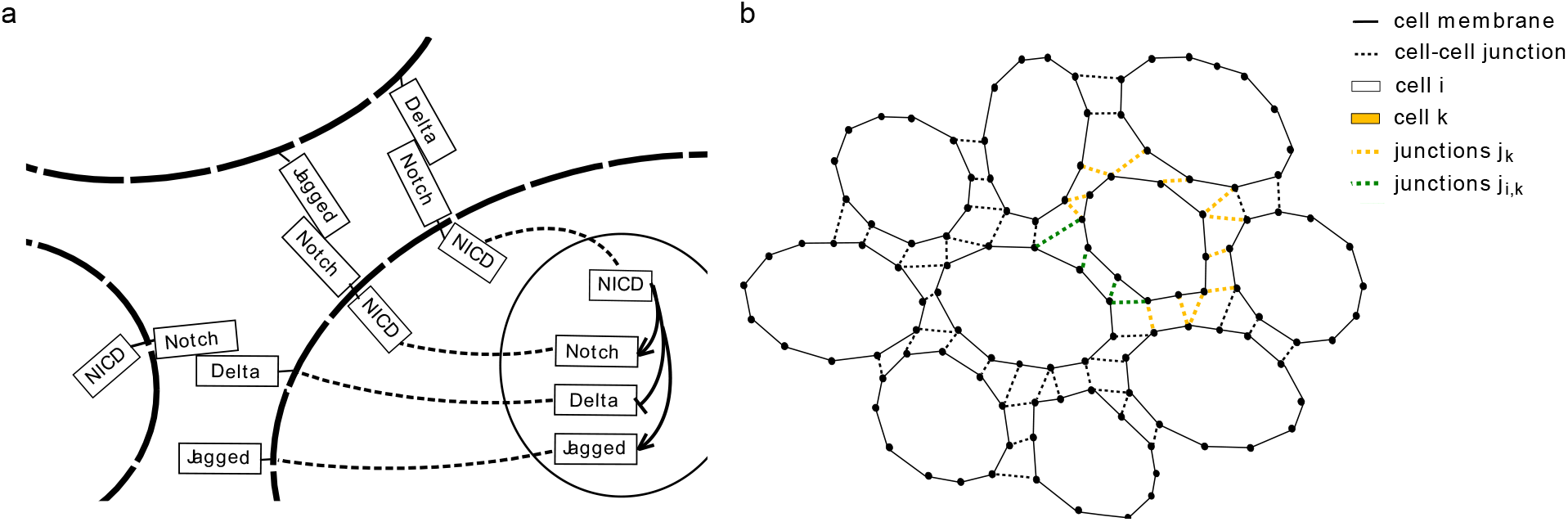
Schematic representation of the Notch pathway and cell representation in LBIBCell. (a) The transmembrane receptor Notch interacts with its transmembrane ligands Delta and Jagged from neighbouring cells. The interaction between the Notch receptor with the Notch ligand leads to the release of the Notch intracellular domain (NICD). NICD works as a transcription factor and modulates the transcription of various genes in the cell nucleus. (b) The amount of Notch receptors and ligands from the cell *i* (blue) available to interact with the cell k (yellow) is dependent on the fraction of shared junctions j_*i,k*_. Consequently, the amount of Notch receptors and ligands available for interaction depends on the cell-cell contact area between the neighbouring cells.

The model represents the dynamics of the Notch receptor (N), the ligands Delta (D) and Jagged (J), and the Notch signal (I), by the following differential equations (Boareto et al, 2015):

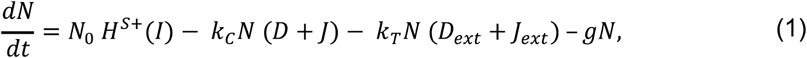

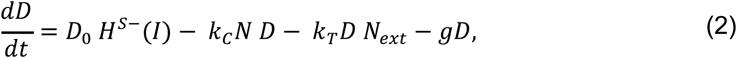

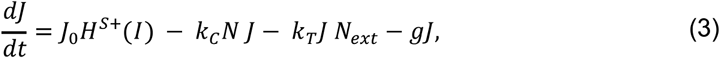

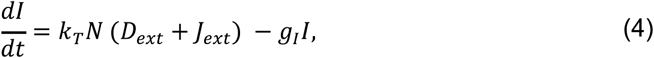

where *g* represents the degradation rate of the proteins, *k_T_* and *k_C_* represent the strengths of *cis*inhibition and *trans*-activation rates respectively; and *N*_0_, *D*_0_ and *J*_0_ are the production rates of Notch, Delta and Jagged, respectively. *N_ext_, D_ext_* and *J_ext_* represent the amount of protein available for binding from the neighbouring cells which is proportional to the shared contact surface (Figure 5b). Shifted Hill functions are considered to represent the effect of NICD (I) on the production rates of Notch receptors and ligands. Shifted Hill functions are defined as *Hs*(*I,λ*) = *H*-(*I*) + *λH*+(*I*) or in simpler notation: *Hs*+(*I*) for activation (λ>1) and *Hs*-(*I*) for repression (λ<1), and λ represents the fold-change in the production rate. The values of the parameters are described in Table 1.

**Table 1.** Parameters of the Notch signalling model (Eqs. 1–4).

| Parameter | Dimension | Value |
| --- | --- | --- |
| $N_0$ | Number of proteins*(Time)<sup>-1</sup> | 6.0 |
| $D_0$ | Number of proteins*(Time)<sup>-1</sup> | 10.0 |
| $J_0$ | Number of proteins*(Time)<sup>-1</sup> | {3.0,...,8.0} |
| $k_C$ | (Time)<sup>-1</sup> | 5.0 10<sup>-6</sup> |
| $k_T$ | (Time)<sup>-1</sup> | 5.0 10<sup>-7</sup> |
| $\gamma$ | (Time)<sup>-1</sup> | 1.0 10<sup>-3</sup> |
| $\gamma_I$ | (Time)<sup>-1</sup> | 5.0 10<sup>-3</sup> |
| $\lambda_N$ | Dimensionless | 2.0 |
| $\lambda_D$ | Dimensionless | 0.0 |
| $\lambda_J$ | Dimensionless | 5.0 |

To model Notch signalling between neighbouring cells that interact on a growing domain, we used `LBIBCell` (Tanaka et al, 2015), an open-source C++ 2D tissue simulation environment (https://bitbucket.org/tanakas/lbibcell/src/master). We combined a set of `BioSolver` (BS) representing cellular processes, and an `ODESolver` (OS) that solves the Notch signalling dynamics.

We used the following basic `BS: Membrane Tension BS, Cell Junction BS, Cell Division BS, Cell Growth BS` and `Cell Apoptosis BS`. This set of BS involves a total of 13 free parameters (Table 2). A detailed description of the software, `BS`, and how we defined a suitable parameter space in `LBIBCell` can be found in (Tanaka et al, 2015; Kokic et al, 2019; Stopka et al, 2019). We set the growth source, such that the default cell cycle time in `LBIBCell` was t_cc_ ~ 2400 LB_time. As pancreatic progenitors divide on average once every 24 hours (Kim et al, 2015), one iteration step in `LBIBCell` corresponds to approximated 36s and the signalling parameters were chosen accordingly (Table 1). We further used the previously developed `Cell Differentiation BS` to represent the cell differentiation process during development (Tanaka et al, 2015). The default cell is of type 1, and cells with a NICD concentration below a certain threshold (100 LB units) change their fate to type 2.

**Table 2.** The parameter space for LBIBCell. The `BS` and their parameters are listed. When the parameter values are set within the specified range, the simulated tissues are epithelial-like.

| BioSolver | Parameter description | Dimension | Valid range [LB] | Value |
| --- | --- | --- | --- | --- |
| **Cell Growth** | Mass added to intracellular grid point | Mass/Time | $\{10^{-3}, \dots, 10^{-4}\}$ | $\{3.5, 7, 15\} 10^{-4}$ |
| | Call frequency | $(\text{Time})^{-1}$ | $\geq 10^{-2}$ | 1 |
| **Cell Membrane Tension** | Hookean spring constant | Force/Length | $\{10^{-3}, \dots, 10^{-2}\}$ | $10^{-2}$ |
| | Call frequency | $(\text{Time})^{-1}$ | $\geq 10^{-2}$ | 1 |
| **Cell-Cell Junction** | Hookean spring constant | Force/Length | $\{10^{-3}, \dots, 10^{-1}\}$ | $5.0 10^{-3}$ |
| | Length of intercellular cell junction | Length | $\{10^{-2}, \dots, 10^0\}$ | $5.0 10^{-1}$ |
| | Radius for junction building | Length | $\{0.5, \dots, 1.5\}$ | 1.0 |
| | Call frequency | $(\text{Time})^{-1}$ | $\geq 10^{-2}$ | 1 |
| **Cell Division Asymmetric** | Cell division area | $(\text{Length})^2$ | $\geq 130$ | 270 |
| | Split factor | Dimensionless | $\geq 2.0$ | $\{2.0, 3.0, 4.0\}$ |
| | Call frequency | $(\text{Time})^{-1}$ | $\leq 10^{-2}$ | $10^{-1}$ |
| **Cell Apoptosis** | Minimal cell area | $(\text{Length})^2$ | $\leq 50$ | 20 |
| | Call frequency | $(\text{Time})^{-1}$ | $\leq 10^{-2}$ | $10^{-1}$ |
| **Neighbor Concentration** | Call frequency | $(\text{Time})^{-1}$ | $\geq 10^{-2}$ | 1 |
| **Cell Differentiation** | Call frequency | $(\text{Time})^{-1}$ | $\geq 10^{-2}$ | 1 |

In addition to the available `BS`, we developed two new `BS`. To model the signalling interactions between neighbouring cells, we developed the Neighbor `Concentration BS`. This BS allows us to retrieve the protein concentrations on the neighbouring cells as well as the fraction of shared junctions. The local protein concentration that is available for interactions is then set to the protein concentration of the neighbouring cell times the fraction of shared junctions. To explore the effects of asymmetric cell divisions, we developed the `Cell Division Asym BS`. The default `Cell Division BS` generates a cell division plane that is perpendicular to the mitotic spindle axis of a cell and intersects it in the middle. Thus, the cell is split symmetrically and daughter cells have approximately the same size. In the new `Cell Division Asym BS`, the intersection point of the division plane can be placed such that the mitotic spindle axis is not split in two halves but in unequal parts, e.g., two thirds and one third. The mother cell is then divided asymmetrically and the daughter cells have different sizes. The intersection point (*x_d_,y_d_*) is defined by the asymmetry factor *a* and is calculated as follows:

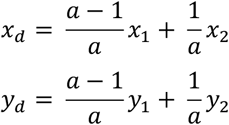

with (*x*_1_,*y*_1_) and (*x*_2_,*y*_2_) being the pair of points that defines the mitotic spindle axis, e.g., the longest axis of a cell. Therefore, for symmetric division *a* = 2 and for asymmetric division *a* > 2. When dividing, cells transmit their own cell fate to their progeny.

## Acknowledgements

This project has received funding from the European Union’s Horizon 2020 research and innovation programme Pan3DP FET Open, EU Horizon 2020 grant agreement No 800981.

## Author contributions

Conceptualization: D.I. and M.B.; Performed research: A.S.; Writing: A.S., M.B. and D.I.; Funding acquisition and project administration: D.I.

